# Binding dynamics of alpha-actinin-4 in dependence of actin cortex tension

**DOI:** 10.1101/2020.03.11.986935

**Authors:** K. Hosseini, Leon Sbosny, Ina Poser, E. Fischer-Friedrich

**Affiliations:** Cluster of Excellence Physics of Life, Technische Universität Dresden, Dresden, Germany; Biotechnology Center, Technische Universität Dresden, Dresden, Germany; Max-Planck-Institut für Zellbiologie und Genetik, Pfotenhauerstraße 108, 01307 Dresden, Germany

## Abstract

Mechano-sensation of cells is an important prerequisite for cellular function, e.g. in the context of cell migration, tissue organisation and morphogenesis. An important mechano-chemical-transducer is the actin cytoskeleton. In fact, previous studies have shown that actin cross-linkers, such as *α*-actinin-4, exhibit mechanosensitive properties in its binding dynamics to actin polymers. However, to date, a quantitative analysis of tension-dependent binding dynamics in live cells is lacking. Here, we present a new technique that allows to quantitatively characterize the dependence of cross-linking lifetime of actin cross-linkers on mechanical tension in the actin cortex of live cells. We use an approach that combines parallel plate confinement of round cells, fluorescence recovery after photo-bleaching, and a mathematical mean-field model of cross-linker binding. We apply our approach to the actin cross-linker *α*-actinin-4 and show that the cross-linking time of *α*-actinin-4 homodimers increases approximately twofold within the cellular range of cortical mechanical tension rendering *α*-actinin-4 a catch bond in physiological tension ranges.

## I. INTRODUCTION

Mechanosensation of cells is pivotal for the regulation of cell adhesion and cytoskeletal force generation. Therefore, it is vital for biological processes such as cell migration, cell shape emergence, tissue organisation and morphogenesis^1–4^. A particularly important role for mechanotransduction inside cells plays the actin cytoskeleton that connects to the extracellular matrix via focal adhesion complexes^2^. Mechanosensation is enabled by mechanosensitive molecules inside cells that transduce mechanical triggers into chemical cues. One way to realize such a mechanochemical transduction is through mechanosensitive bonds: protein-protein bonds subject to mechanical tension may either strengthen (catch bonds) or weaken (slip bonds) in response to tensile mechanical stress. In turn, catch (slip) bonds enhance (decrease) their bond lifetime in response to tension^5^. Furthermore, catch (slip) bond dynamics is associated with an increased (decreased) fraction of bound proteins in the presence of mechanical tension.

Previous studies put forward that the actin cross-linker *α*-actinin-4 forms catch bonds: Yao *et al.* showed stress-enhanced gelation of *in vitro* actin networks cross-linked by *α*-actinin-4 indicating catch bond behaviour^6^. Previous studies with *Dictyostelium* cells demonstrated slowed *α*-actinin-4 turnover in the actin cytoskeleton in response to compressive stress^7^ and accumulation of *α*-actinin-4 in response to suction pressure from micropipette aspiration^8^. However, to date, there is no study that showed a quantitative relation between *α*-actinin-4 cross-linking lifetimes and mechanical tension in the cytoskeleton. The goal of our study is to close this gap by establishing a new technique that combines parallel plate assays of non-adherent cells based on atomic force microscopy (AFM) with photo-bleaching of fluorescently labeled *α*-actinin-4 and mathematical modelling of cross-linker binding kinetics.

As many actin cross-linkers, *α*-actinin-4 binds at a high affinity to a dumbbell-shaped antiparallel homodimer in the cytoplasm^9^, where each monomer inside the homodimer provides and F-actin-binding domain (ABD), which allows *α*-actinin-4 homodimers to serve as cross-linkers in actin cytoskeletal networks. Previous studies have characterized the biochemical features of *α*-actinin-4 actin-binding: each *α*-actinin-4 monomer inside the homodimer contains two N-terminal calponin homology (CH) domains which are normally tethered in a closed conformation to form the ABD^9,10^. An additional actin binding site outside the ABD of *α*-actinin-4 is present but normally buried inside the molecule in the closed conformation. Alternatively, an open conformation can be adopted by *α*-actinin-4 with a higher actin-binding affinity: this can be achieved through biochemical modifications such as phosphorylation at Y265^10^. The exposure of this additional actin binding site through mechanical stretching of the cross-linking homodimer has been suggested to be at the heart of mechanosensitive binding dynamics of *α*-actinin-4^10^. Furthermore, the binding affinity of normal *α*-actinin-4 is regulated by *Ca*^2+^ ions through a C-terminal calmodulin-like *Ca*^2+^-binding domain where the presence of *Ca*^2+^ reduces the binding affinity of *α*-actinin-4 to actin^9,10^.

In our study, we quantiatively characterize the lifetime of fluorescently labeled *α*-actinin-4 constructs at the actin cortex in HeLa cells in mitotic arrest. To this end, we combine confocal imaging of the cortex and fluorescence recovery after photo-bleaching (FRAP) with a parallel plate confinement assay of cells that allows to measure the momentary mean-field mechanical tension in the actin cortex as described previously^11–13^. We use our results on FRAP-derived cortical residence time at given values of cortical tension to infer the cross-linking lifetime of *α*-actinin-4 in the cortex using a simple mathematical model of cross-linker binding as has been previoulsy suggested by^14^. We extend this model such that it includes the dependence of cross-linking lifetimes on mechanical tension in the actin cortex and fit model predictions to experimental data. Furthermore, we present passive and active methods to manipulate cortical tension in individual cells and monitor cross-linker concentration concomitantly.

## II. MATHEMATICAL MODEL OF TENSION-DEPENDENT CROSS-LINKER BINDING DYNAMICS

We adapt a coarse-grained description of cross-linker dynamics developed by Mulla et al.^14^ that allows to connect the ensemble-averaged lifetime of a cross-linking protein at the cortex with the lifetime of its cross-linking state. We formally extend the model to include the change of cross-linking lifetimes under mechanical loading.

The model includes the dynamic exchange of three populations of cross-linker protein homodimers: i) a well-stirred cytoplasmic population *c_cyt_*(*t*), ii) homodimers bound at the cortex with a single ABD *c_sb_*(x, *t*) (experiencing no mechanical loading), and iii) homodimers bound with two ABDs *c_cl_*(x, *t*) (cross-linking state), see Fig. 1. Populations *c_sb_*(x, *t*) and *c_cl_*(x,*t*) are bound to the cortex and contribute to the fluorescence signal during FRAP measurements. The defining equations of homodimer dynamics are

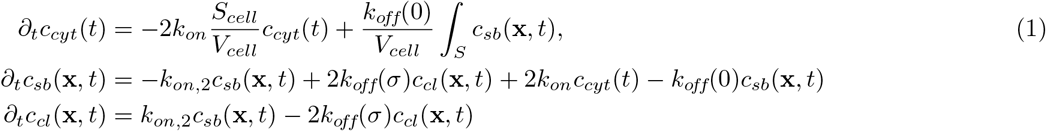

where *S_cell_* and *V_cell_* are the cell surface area and the cell volume, respectively, and *∫_S_ c_sb_* denotes the integration of *c_sb_* over the cell surface. The attachment rate *k_on_* characterises the binding rate of an ABD inside cytosolic homodimers to the actin cortex. The attachment rate *k*_*on*,2_ characterises the rate of actin-binding of the second ABD of a homodimer that is already bound via one ABD. Furthermore, the rate *k_off_*(*σ*) describes the average rate of unbinding of one ABD of a cross-linking homodimer at a mean-field cortical tension of *σ*. At steady state, the fraction of cortical proteins *F_cor_*(*σ*) = *N_cor_*/*N_tot_* takes the following form

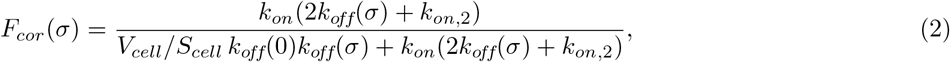

where *N_cor_* = (*c_cl_* + *c_sb_*)*S_cell_* is the number of cortex-associated cross-linkers, *N_tot_* = *N_cor_* + *N_cyt_* is the overall number of cross-linker proteins and *N_cyt_* = *c_cyt_V_cell_* is the number of cytoplasmic proteins. The ratio *R*(*σ*) = *N_cor_/N_cyt_* of cortical to cytoplasmic homodimers is given by

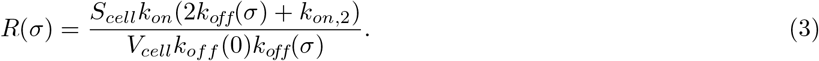

**Figure 1.**
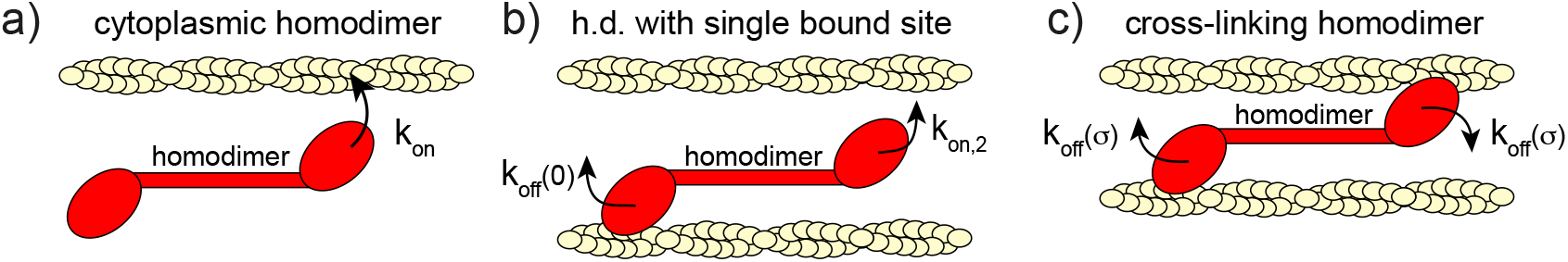
Schematic of actin cross-linker binding dynamics according to our model. a) Cytoplasmic (well-stirred) homodimers bind with a rate *k_on_* to actin filaments. b) Homodimers bound via a single ABD either detach from actin filaments at a rate *k_off_*(0) or attach also via the second ABD with a rate *k*_*on*,2_. c) Crosslinking homodimers may detach at either binding site with a rate *k_off_* (*σ*), where *σ* is the mean-field mechanical tension in the actin cortex.

In the sequel of the manuscript, we will assume a simple catch bond behavior of the cross-linker with a linear tensiondependence of cross-linking time *k_off_*(*σ*)^−1^ = (1+*α σ*)*k_off_*(0)^−1^. This approach can be considered as a Taylor expansion up to first order of the tension dependence of cross-linker lifetime.

From Eqn. (2), we determined the average residence time of a homodimer at the cortex by calculating the time evolution of the probability of a freshly bound homodimer to remain in a single-bound state or a cross-linking state at the cortex. Let *p_sb_*(*t*) and *p_cl_*(*t*) denote these probabilities with *p_sb_*(0) = 1 and *p_cl_* (0) = 0. Then, they evolve in time according to

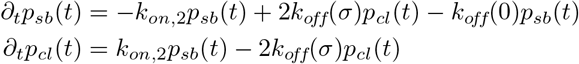

This set of equations can be solved analytically as

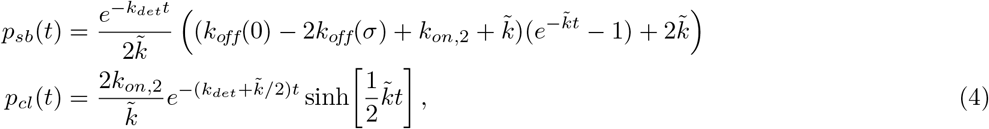

where

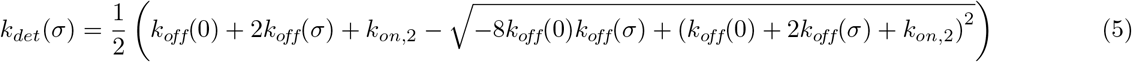

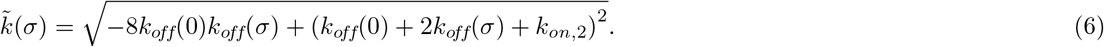

Solutions for differrent parameter sets are presented in Fig. 2a-c, left column. The dynamics exhibits a slowest time scale *τ_det_* = *k_det_*(*σ*)^−1^. This time scale constitutes an approximation to the residence time at the cortex and characterizes the slowest time scale in the recovery after photo-bleaching (see Fig. 2a-c, middle column, black dashed lines). In the following, we will refer to *k_det_*(*σ*)^−1^ as cortical residence time. In fact, the cortical dynamics can be approximated by a simple reaction kinetics of the kind

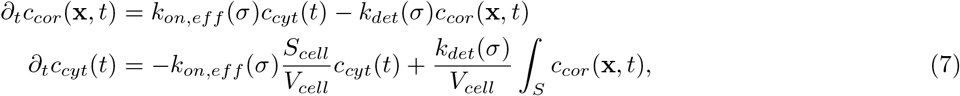

where *c_cor_* captures the concentration of all homodimers at the cortex (regardless of single-bound or cross-linking). Here, the condition of equal steady state concentrations at the cortex and in the cytoplasm is used to determine the effective attachment rate

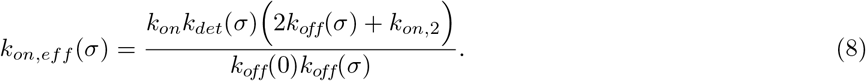

**Figure 2.**
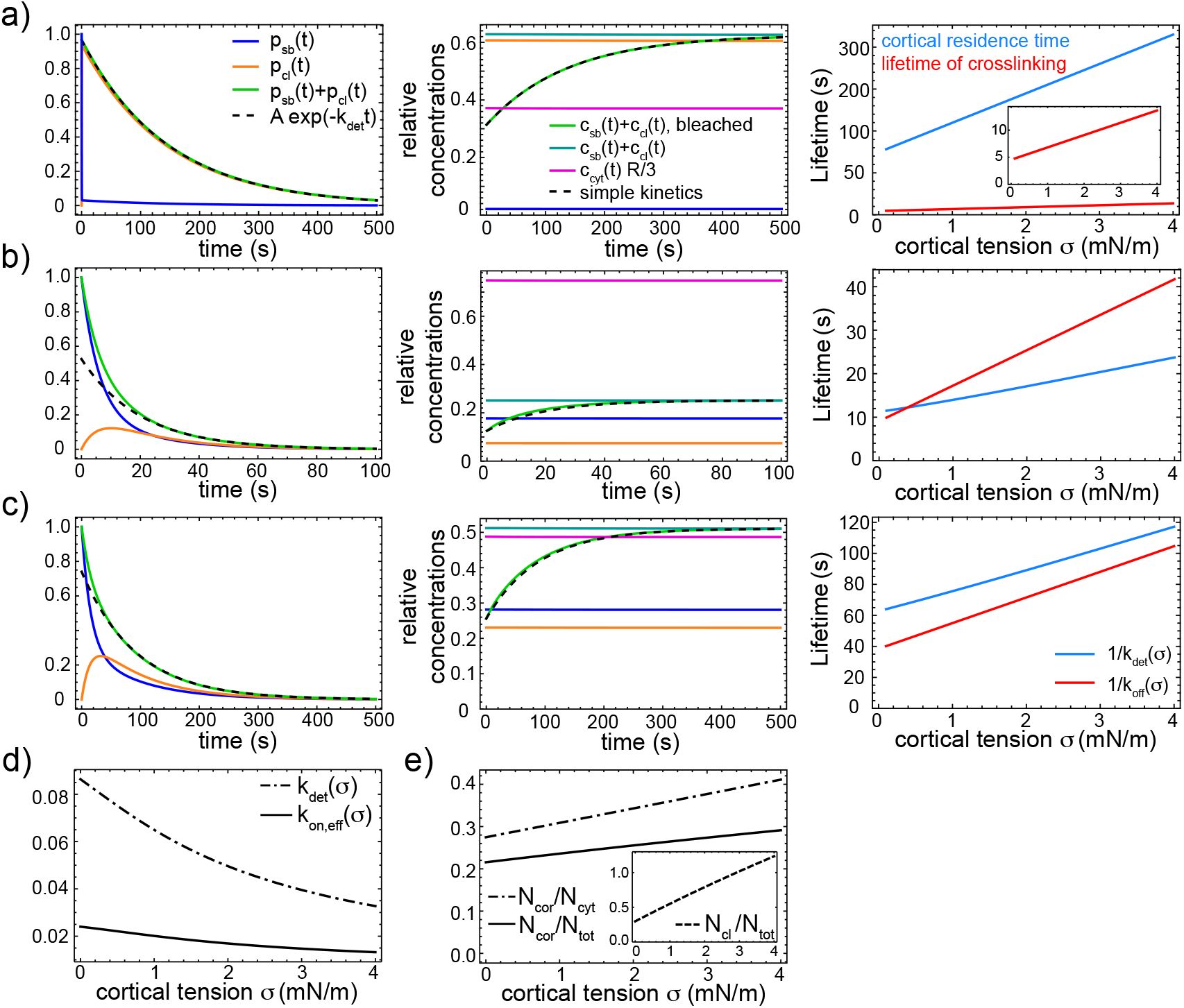
a-c) Coarse-grained model of homodimer dynamics for three different parameter sets. *Left panel column:* Time evolution of the probability of a homodimer to be in a cross-linked (*p_cl_*, orange curve) or a single-bound state (*p_sb_*, blue curve) according to Eqn. (4). The event of initial cortical attachment is assumed to be at t=0 s, i.e. *p_sb_*(0) = 1 and *p_cl_*(0) = 0. The associated exponential decay with the slowest time scale (1 /*k_det_*(*σ*)) of the model dynamics is shown by the dashed line. *Central panel column:* Simulation of FRAP recovery according to model Eqn. (2) after bleaching of a small region of the cortex (green: concentration at the cortex in the bleached region, stationary state concentrations: cytoplasmic scaled (pink), total cortical (turquoise), cortical single-bound (blue), cortical cross-linking (orange)). The corresponding simplified kinetics as described by Eqn. (7) is shown by dashed black lines. *Right panel column:* Cross-linking lifetime (red) and predicted cortical residence time (blue) of homodimers in dependence of cortical tension *σ*. The total residence time of the homodimer at the cortex follows qualitatively the same trend as the cross-linking lifetime. d) Attachment and detachment rate of the approximating simple kinetics in dependence of cortical tension. e) The fraction of cortex-associated homodimers increases with tension in the scenario of a simple catch bond behaviour due to increased cross-linking times. Inset: fraction of cross-linking homodimers in dependence of tension. Model parameters for (d) and (e) are the same as in (b). Parameters used in simulations corresponding to (a) were *R* = 10*μ*m, *k_on_* = 0.021*μ*m, *k*_*on*,2_ = 6.6 s^−1^, *k_off_*(0) = 0.22 s^−1^, *α* = 0.5 × 10^3^*m/N*, *σ* = 2 mN/m and *k_off_*(*σ*) = *k_off_* (0)/(1 + *ασ*). Parameters corresponding to plots in (b) and (c) were R = 10*μ*m, *σ* = 2 mN/m and otherwise taken from Table I, second and fourth row, respectively. In all FRAP simulations, the bleached percentage of the cortex was chosen to be 1%.

The recovery dynamics of the corresponding simplified reaction kinetics after bleaching is illustrated in Fig. 2a-c, central panels (black dashed lines). It approximates the full homodimer dynamics closely if either the population of single-bound homodimers or of cross-linking homodimers dominates at the cortex. The latter scenario is realized in the model dynamics in Fig. 2a. In general, our model predicts that *k_det_*(*σ*)^−1^ captures the slowest time scale in recovery of cortical *α*-actinin-4 fluorescence after photo-bleaching. It is noteworthy that both, attachment and detachment rate of the simplified reaction kinetics, depend now on cortical tension (Fig. 2d), which is in contrast to the full model.

Our analysis shows that a catch or slip bond behavior of a cytoskeletal homodimer is qualitatively reflected in the tension-dependence of cortical residence time of the homodimer (see Fig. 2, rightmost panels). However, cortical residence times *k_det_*(*σ*)^−1^ can be smaller or larger than cross-linking lifetimes *k_off_*(*σ*)^−1^. Further, the slopes of their respective tension-dependences may be distinct from each other (see Fig. 2a-c, right panels). The cortical residence time strongly exceeds the cross-linking lifetime if the fraction of cross-linking homodimers is substantial (see Fig. 2a,b, right panels). In this case, homodimers undergo on average several cycles of rebinding from the single-bound to the cross-linking state before they go back into the cytoplasm. By contrast, the average cortical residence time may also be smaller than the average cross-linking lifetime (see Fig. 2b, rightmost panel). In this case, we have a substantial fraction of single-bound homodimers at the cortex that never go into the cross-linking state but instead drop off quickly into the cytoplasm thereby bringing down the average cortical residence time. Corresponding to an increased cortical residence time, our model predicts an increased cortex-bound fraction of homodimers at higher cortical tension in the case of catch bond dynamics (see Fig. 2e).

It is noteworthy that all four model parameters (*k_on_*, *k*_*on*,2_, *k_off_* (0) and *α*) can be determined if effective detachment rate *k_det_* and fraction of cortically bound homodimers *F_cor_* have been determined for two different known values of cortical tension *σ*_1_ and *σ*_2_. In this case, the four expressions for *F_cor_*(*σ*_1_), *F_cor_*(*σ*_2_), *k_det_*(*σ*_1_) and *k_det_*(*σ*_2_) as given by Eqn. (2) and (5), respectively, can be numerically inverted to obtain the four unknown model parameters.

## III. EXPERIMENTAL RESULTS

We now want to use our theoretical insight to extract the cross-linking lifetime of *α*-actinin-4 from the cortical residence time in the actin cortex of live cells. As a cellular model system, we utilized HeLa cells expressing a fluorescently labeled *α*-actinin-4 construct (see Materials and Methods). Measured cells were arrested in mitosis to take advantage of the adoption of a well-defined cell-cycle stage and of a round deadhered cell shape with a largely uniform actin cortex. Further, we and others reported a particularly high active cortical tension in mitosis^11,15,16^ which allows to explore a large cortex tension range in combination with tension-inhibiting drugs. At the beginning of the experiment, cells cultured in a glass-bottom petri dish were mounted on a confocal microscope in combination with an atomic force microscope (Figure 3a, Materials and Methods). We used this setup to quantify the cortical residence time of fluroescent *α*-actinin-4 by photo-bleaching and imaging of subsequent fluorescence recovery at the cortex (Figure 3b, Materials and Methods). A characteristic time scale *τ* of the recovery was determined by a fit of the recovering fluorescence intensity with a simple exponential function *A*(1 – exp(-*t/τ*)) + *B*. This time scale was identified with the cortical residence time of *α*-actinin-4 at the cortex (Materials and Methods). During FRAP measurements, the cell under consideration was confined via the wedged cantilever of an atomic force microscope. In this way, a time series of the AFM force and the cell height was recorded (Materials and Methods)^11,12^. An exemplary readout is shown in Figure 3c. We previously showed that large cell surface tension in mitosis drives cells into droplet shapes with constant mean curvature in mechanical confinement. Using the previously developed analysis scheme for such droplet-shaped cells, we could determine cortical tension of measured cells from the corresponding AFM readout and cellular imaging^11,12,17,18^.

**Figure 3.**
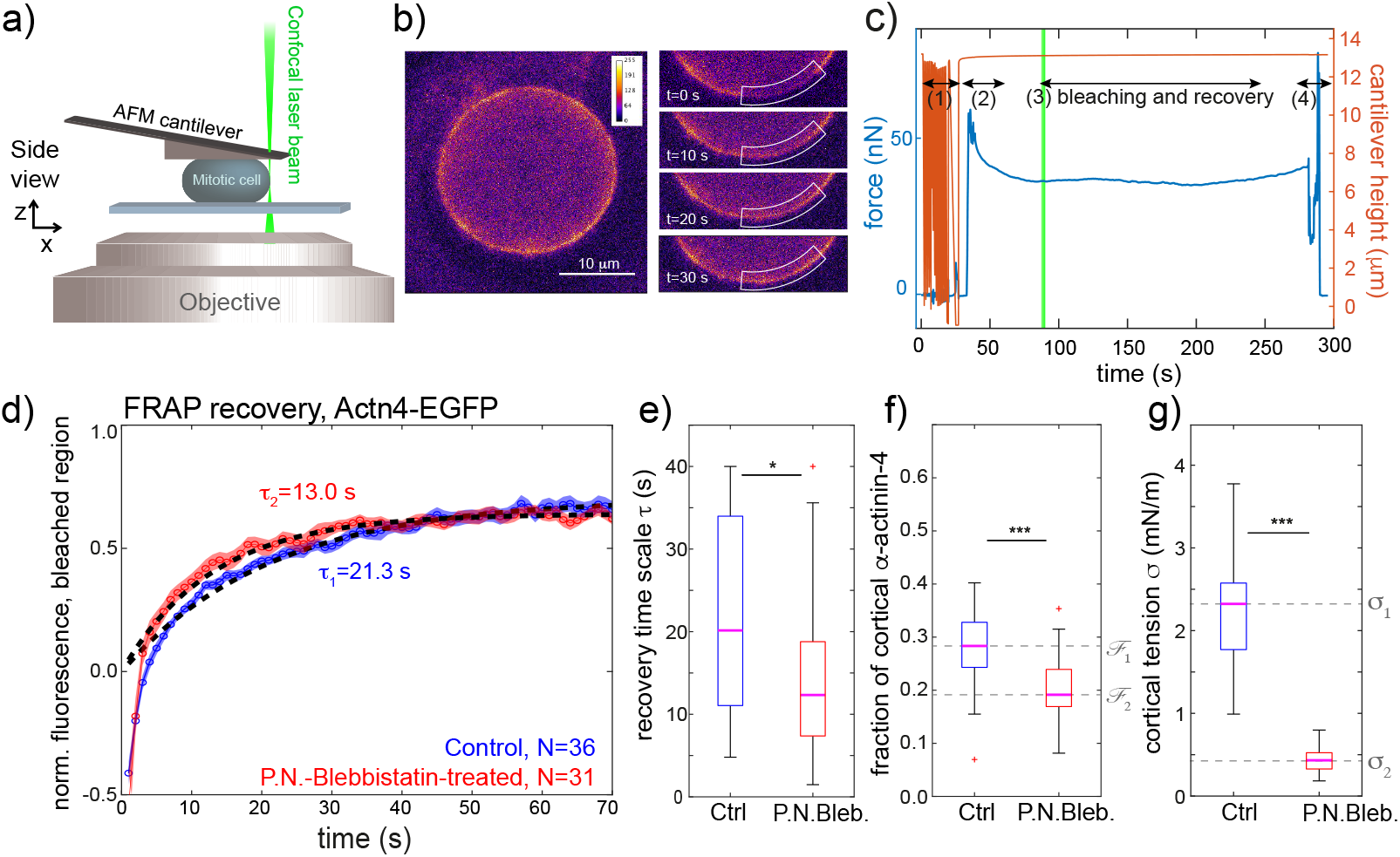
Photo-bleaching of *α*-actinin-4 in the mitotic cortex in combination with an AFM confinement assay. a) Schematic of the experimental setup. Confocal imaging and photo-bleaching of cells in mitotic arrest was combined with parallel plate confinement by wedged AFM cantilevers. Confocal imaging was performed in the equatorial plane of the cell (maximum crosssectional area). The AFM readout was the confinement height and the confinement force of the cell over time. b) Exemplary confocal image of the equatorial plane of a mitotic cell (left panel) and fluorescence evolution after photo-bleaching (panels on the right). The bleached region was indicated by a white boundary. c) Exemplary force and cantilever height readout during the experiment. In phase (1), the cantilever is approached on the glass substrate of the cell culture dish. In phase (2), the retracted cantilever is moved over the cell. At the beginning of phase (3), photo-bleaching is performed and the cortex is imaged during fluorescence recovery. In phase (4), the cantilever is removed from the cell and lifted. d-g) Experimental results from the combined assay on HeLa cells expressing Actn4-EGFP. d) Averaged recovery curves of normalized fluorescence intensity in control conditions (N=36, blue curve) and of cells with pharmacologically reduced cortical tension (N=31, red curve, incubated with Para-Nitro-Blebbistatin at 10 *μ*M). Red and blue shaded regions indicate the respective standard error of the mean for each condition. Averaged normalized curves were fitted with an exponential increase (black dashed lines), with fit time scales of *τ* = 21.3s (control) and 13.0s (P.N.Bleb., 10*μ*M). e) Recovery times from fits of individual recovery curves in control and Blebbistatin-treated conditions corresponding to a. f) Corresponding estimated fractions of cortical *α*-actinin-4. g) Corresponding cortical tensions in control and Blebbistatin-treated conditions. (Measurements are representative for at least two independent experiments. ^*ns*^*p* > 0.05, \**p* < 0.05, \*\**p* < 0.01, \*\*\**p* < 0.001)

We first investigated a HeLa cell line with GFP-labeled *α*-actinin-4 (Actn4-EGFP). Corresponding averaged normalised fluorescence recovery curves are presented in Fig. 3d (blue curve: control conditions, N=36, red curve: tension-reduced condition, N=31, Para-Nitro-Blebbistatin 10 *μ*M). Fluorescence intensities were normalised before averaging as described in Materials and Methods. Fitting an exponential function, we find recovery times of *τ*_1_ = 21.3 s and *τ*_2_ = 13 s in control and tension-reduced conditions, respectively. In addition, we fit the recovery curve for each cell individually and find median recovery time scales that are in close correspondence with time scales obtained from averaged recoveries (Fig. 3e). Median values of active cortical tension in control and tension-reduced conditions were *σ*_1_ = 2.3 mN/m and *σ*_2_ = 0.43 mN/m, respectively (Fig. 3g). According to our model, cortical recovery times *τ* correspond to *k_det_* (*σ*)^−1^. Thus, the model predicts an increased cortical residence time at higher tension (Fig. 2a-c, right panel). This prediction is confirmed by our experimental observations (Fig. 3e). Further in agreement with our model predictions, we measured decreased actin binding in tension-reduced conditions: we found median values of the fraction of cortical *α*-actinin-4 to be 0.28 in control conditions and 0.19 in tension-reduced conditions, respectively (Fig. 3f).

To further test how mechanical tension influences the actin binding dynamics of *α*-actinin-4, we modified mechanical tension over time in the actin cortex of individual cells while jointly monitoring the fraction of cortex-bound protein. According to our model, a catch bond dymamics of *α*-actinin-4 would give rise to an increase of cortex-bound protein when cortical tension increases (Fig. 2d). We manipulated cortical tension in individual cells in two different ways: i) by inducing additional cortical tension via mechanically dilating the actin cortex through external perturbation (Fig. 4), or ii) by pharmacologically increasing active mechanical tension in the actin cortex by photo-deactivation of the myosin inhibitor Blebbistatin (Fig. 5).

**Figure 4.**
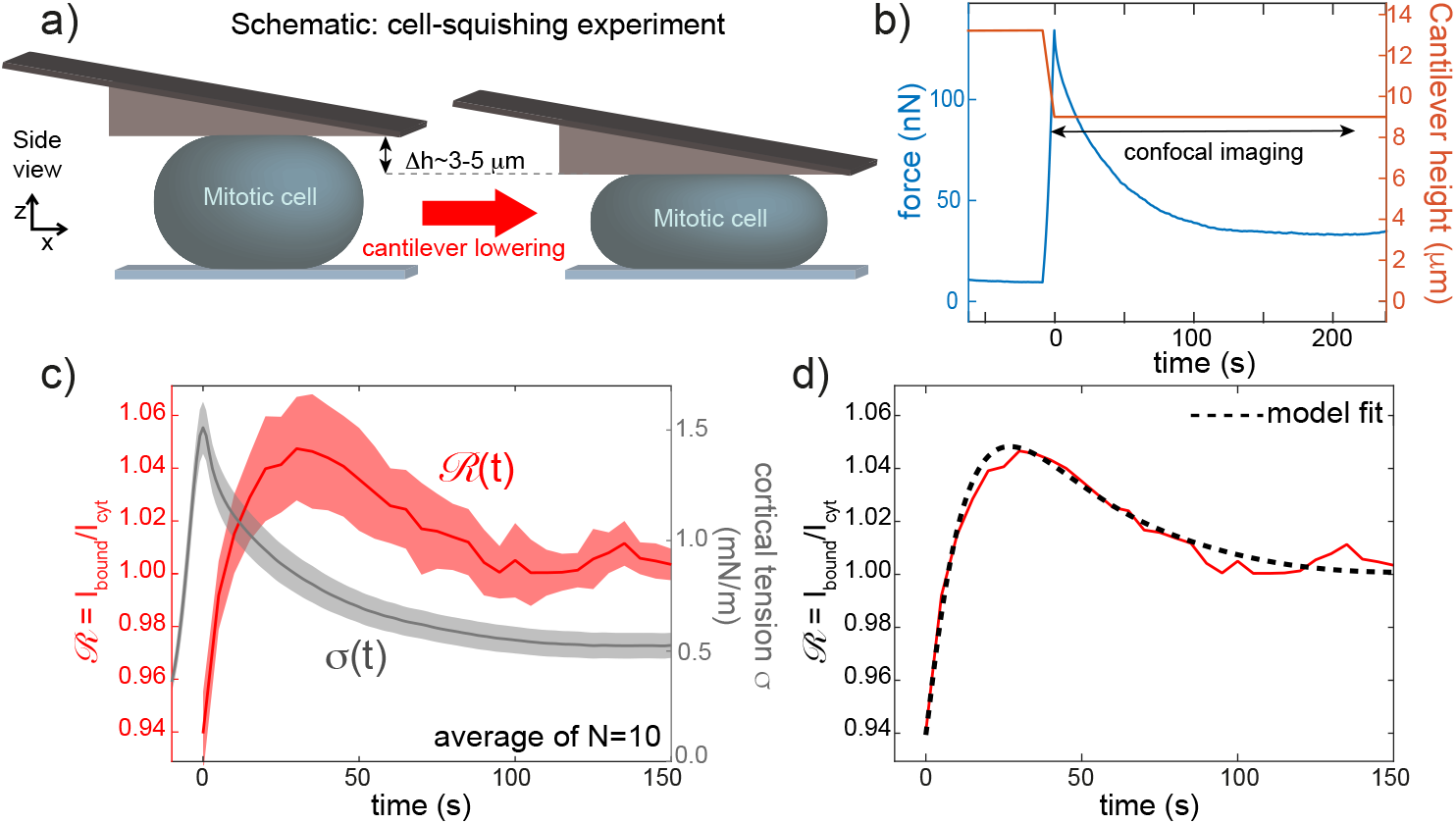
Cell-squishing experiments: fluorescence of *α*-actinin-4 (Actn4-EGFP) increases at the cortex after transient mechanically-induced increase in actin cortex tension. a) Schematic of the experimental procedure. Cells were mechanically perturbed by a fast lowering of the AFM cantilever onto the cell. Thereby, a mechanical stretch of cortical area was induced. Cells were in mitotic arrest and treated with the ROCK inhibitor Y27632 (5 *μ*M). ROCK inhibition was necessary to avoid blebbing during mechanical cell squishing. b) Exemplary force and cantilever height readout during the experiment. While the cantilever is lowered, cellular confinement force increases. Afterwards, the confinement force relaxes to a new steady state force. c) Averaged time evolution of cortical tension and normalised *α*-actinin-4 fluorescence at the cortex during and after mechanical squishing of the cell. Averages were taken over ten cells. Mechanical tension peaks right after cantilever lowering has been stopped. Then, tension drops due to viscoelastic relaxation of deformation-induced cortical mechanical tension^12^. Concomitantly, we observe a peak of *α*-actinin-4 fluorescence at the cortex at ≈ 30 s shortly after the peak in mechanical tension at the actin cortex. Error bars indicate standard errors of the mean. d) Our model on protein dynamics provides an excellent fit (dashed black line) to the measured time evolution of cortical *α*-actinin-4 fluorescence (as depicted in c). Fitting parameters are given in Table I, second row. According to our model, the transient peak in cortical intensity at *t* ≈ 30 s stems from the tension increase and the resulting increase in cross-linking lifetime *k_off_*(*σ*)^−1^. The elevated stationary state value at long times can be accounted for by an increase of the surface to volume ratio due to increased cell confinement.

We will first discuss the effect of increase of cortical tension by cortex dilation through external perturbation. Cortical dilation was achieved by applying a step of cell-squishing using the wedged cantilever of an AFM (Fig. 4a, Materials and Methods). We showed previously that confined mitotic cells adopt droplet shapes at a cell volume that is largely independent of cell height^11^. Consequently, cell surface area must increase if cells are squished. Compression steps in our experiment reduced cell height by 3 – 5 *μ*m at a speed of 0.5 *μ*m/s. The corresponding cell surface increase can be estimated to be between 6.5 – 15% at typical cell volumes^11^. To avoid cell blebbing upon cell-squishing, cells were treated with the ROCK inhibitor Y27632 (5 *μ*M) which leads to a reduction of myosin activity and slowed viscoelastic relaxation of stresses after cortex dilation^12,19^. During cantilever lowering, the measured AFM force quickly rises. Once the cantilever comes to a halt, the force relaxes to a new steady state within a time interval of ≈ 100 s indicating the viscoelastic relaxation of deformation-induced cortical mechanical tension over time (Fig. 4b). We monitored the time evolution of the intensity ratio of cortical to cytoplasmic fluorescence starting from the moment where the cantilever comes to a halt until mechanical tension has been relaxed. We calculated averaged values of normalised fluorescence ratio for ten cells (red curve, Fig. 4c) jointly with averaged tension evolution (grey curve, Fig. 4c). We see an intermediate peak in the averaged cortex-to-cytoplasm ratio of fluorescence at a time *t* ≈ 30 s after the uniaxial compression step (see Materials and Methods). This peak is followed by a relaxation to a new elevated steady state value. A fit of our model that conforms with measured FRAP recovery times and cortical fractions (see Fig. 3d-f) can account for our observation (Fig. 4d, black dashed line, model parameters given in Table I, second row); the intermediate concentration rise stems from the transient peak in cortical tension and a corresponding transient dip in detachment rates *k_off_*(*σ*). Furthermore, the elevated final steady state originates from an increase in the surface to volume ratio during cell-squishing and the adoption of a corresponding new equilibrium state of the dynamic system given by Eq. (2).

**Table I.** Estimated model parameters for the binding of fluorescently-labeled *α*-actinin-4 homodimers for two different cell lines expressing either Actn4-EGFP or Actn4-mKate2. For both cell lines, parameters have been estimated according to the simple (green) or elaborate (brown) analysis pathway (see Fig. 6 and Materials and Methods). Here, 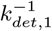 and 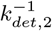 denote cortical residence times at tension values of *σ*_1_ (control conditions) or *σ*_2_ (tension-reduced conditions). Cortical tension values *σ*_1_ and *σ*_2_ were chosen as median tension values from Fig. 3g and k as well as Fig. 5d in respective measurement conditions. Error ranges are obtained by error propagation (see Materials and Methods). Parameter values of the brown analysis pathway have to be considered as more accurate.

| Cell line | $k_{on} (\mu\text{m/s})$ | $k_{on,2} (1/\text{s})$ | $k_{off}(0) (1/\text{s})$ | $\alpha (10^3 \text{m/N})$ | $k_{det,1}^{-1} (\text{s})$ | $k_{det,2}^{-1} (\text{s})$ | $F_{cor,1}$ | $F_{cor,2}$ |
| --- | --- | --- | --- | --- | --- | --- | --- | --- |
| Actn4-EGFP | 0.036 ± 0.013 | 1.61 ± 2.61 | 0.34 ± 0.20 | 0.55 ± 0.22 | 21.6 | 13 | 0.19 | 0.28 |
| Actn4-EGFP | 0.043 ± 0.003 | 0.035 ± 0.009 | 0.11 ± 0.0034 | 0.9 ± 0.19 | 21.6 | 13 | 0.25 | 0.22 |
| Actn4-mKate2 | 0.026 ± 0.004 | 0.009 ± 0.025 | 0.022 ± 0.012 | 0.92 ± 0.43 | 101.7 | 64.1 | 0.51 | 0.46 |
| Actn4-mKate2 | 0.025 ± 0.002 | 0.026 ± 0.01 | 0.027 ± 0.002 | 0.41 ± 0.18 | 101.7 | 64.1 | 0.51 | 0.46 |

As an alternative method to test the tension dependence of *α*-actinin-4 in single cells, we used the possibility to increase active cortical tension in the cortex of single cells through optical manipulation of the myosin inhibitor Blebbistatin^11,12,16^. When exposed to blue light, Blebbistatin becomes inactive and ceases to reduce myosin activity which gives rise to a boost in myosin activity and cortical tension^12,20^. In the following, we will refer to this effect as indirect photo-activation of myosin. To employ indirect photo-activation, we generated a HeLa cell line expressing an mKate2-labeled construct of Actn4 (Materials and Methods) whose fluorescence excitation is not in the range of blue light^21^. In turn, for this cell line, fluorescent imaging of Actn4-mKate2 does not affect Blebbistatin activity^22^. We first characterized cortical residence time of mKate2-labeled *α*-actinin-4 by FRAP measurements (Fig. 5). We find qualitatively analogous results to the GFP-labeled cell line, i.e. cortical residence times and affinity to cortical actin are larger at conditions of high cortical tension (Fig. 5a-d). Interestingly, we observe significantly longer cortical residence times than for the GFP-labeled construct. We speculate that C-terminal tagging of *α*-actinin-4 might interfere with C-terminally located *Ca*^2+^-binding. This could in turn modify the actin affinity of the *α*-actinin-4 homodimer in a fluorophore-dependent manner^9^.

**Figure 5.**
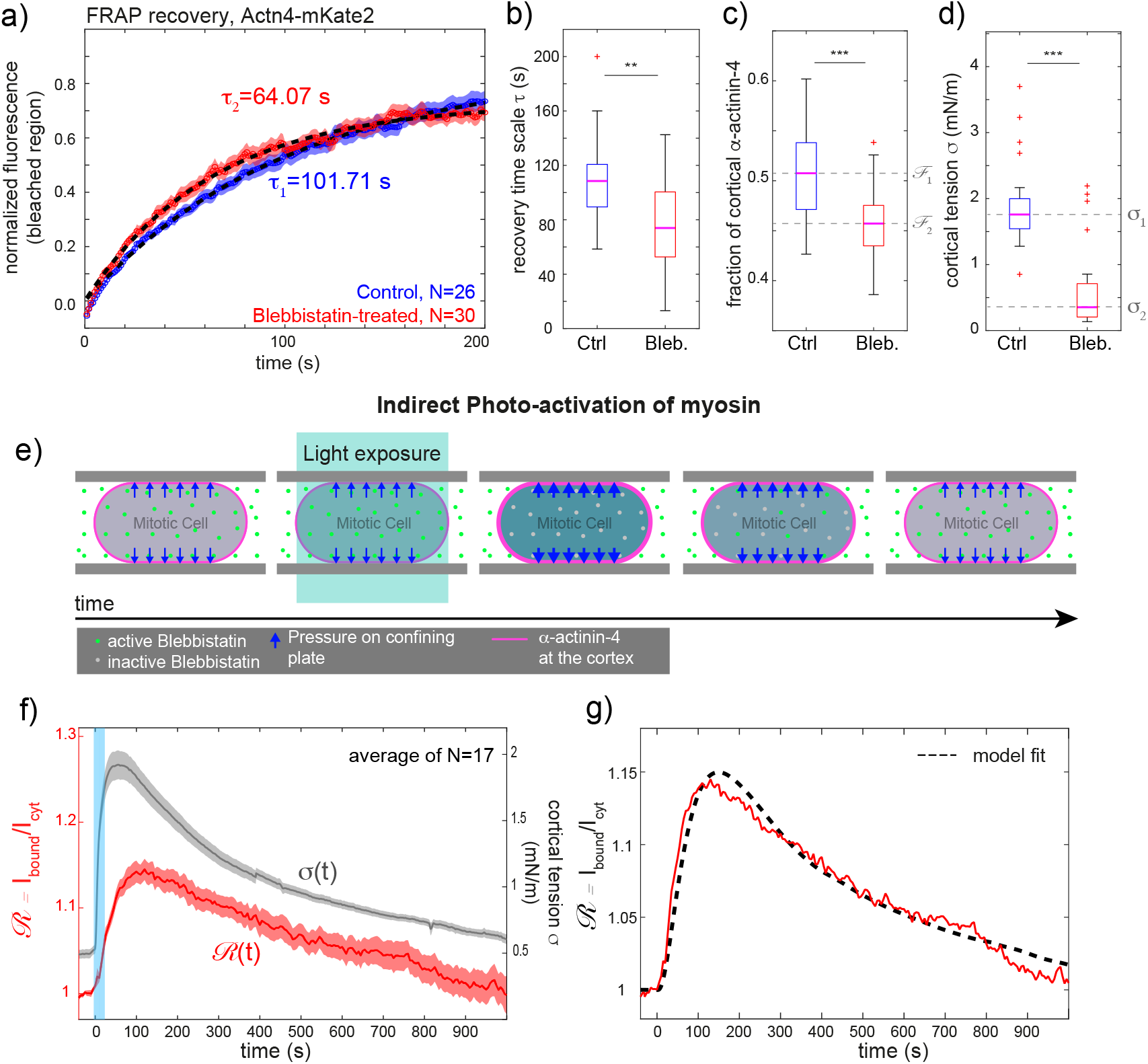
Experimental results from FRAP and indirect photo-activation of myosin with red-fluorescently-labeled *α*-actinin-4 (Actn4-mKate2) in mitotic HeLa cells. a) Averaged recovery curves of *α*-actinin-4-mKate2 after photo-bleaching at the mitotic cortex: control (N=26, blue curve), tension-reduced (N=30, red curve, incubated with Blebbistatin at 10 *μ*M). Red and blue shaded regions indicate the respective standard error of the mean for each condition. Fluorescence intensities were normalized. Averaged curves were fitted with an exponential recovery (black dashed lines), with fit time scales of *τ* = 101.71s (control) and 64.07 s (Bleb., 10*μ*M). b) Recovery times from fits of individual recovery curves in control and Blebbistatin-treated conditions corresponding to a. On average, recovery is quicker if cells were treated with Blebbistatin. Median values are 108.5 s (control) and 72.8s (Blebbistatin at 10*μ*M). c) Estimated fraction of cortical *α*-actinin-4 in control and Blebbistatin-treated conditions corresponding to a. Median values of fit recovery time scales are 0.51 (control) and 0.46. d) Cortical tensions in control and Blebbistatin-treated conditions corresponding to measurements presented in a. e) Schematic of cellular response upon indirect photo-activation of myosin. f) Time evolution of the cortex-to-cytoplasm ratio 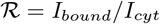 of Actn4-mKate2 fluorescence (red curve) and corresponding time evolution of cortical tension (grey curve). The blue-shaded region indicates the time interval during which cells were scanned with blue laser light (488 nm) to inactivate Blebbistatin inside the cell. g) Averaged time evolution of cortex-to-cytoplasm ratio 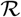 (red curve, as in panel e) with corresponding model fit (black dashed line). Fitting parameters are given in Table I, 4th row. Curves were averaged over 17 cells. Before averaging, intensity curves were normalised to take the value one before photo-inactivation. (Measurements are representative for at least two independent experiments. ^*ns*^*p* > 0.05, \**p* < 0.05, \*\**p* < 0.01, \*\*\**p* < 0.001)

In a next step, we carried out experiments, where cortical tension was dynamically varied by indirect photo-activation of myosin. To this end, we mechanically confined mitotic cells with the AFM cantilever in the presence of a low dose of Blebbistatin in the medium (2.5μM). During steady confinement, we started a time series of confocal imaging of the equatorial cross-section of the cell accompanied by a time series readout of AFM force and cantilever height. After recording 5-10 intial frames in a state of inhibited myosin and reduced cortical tension, we laser-scanned the cell for ≈ 10 s with blue light (see Materials and Methods) and then continued the time series of confocal imaging. The blue light exposure leads to a quick rise of AFM force due to inactivation of Blebbistatin in the cell reflecting an increase of cortical tension by ~ 4-fold (Fig. 5e,f). Over time, active Blebbistatin reenters the cell and cortical tension slowly drops over a time span of 10 – 20 min. We performed an image analysis of the accompanying confocal time series of the cortical and cytoplasmic *α*-actinin-4 population and calculated the cortex-to-cytoplasm ratio of fluorescence intensity (Fig. 5f, red curve, see Materials and Methods). We find on average a trend of a delayed rise of this intensity ratio, indicating a transient increase of *α*-actinin-4 actin binding affinity. Our model can account for this observation in terms of the transient tension rise of cortical tension in combination with a catch bond binding of *α*-actinin-4. A corresponding fit of predicted model dynamics is shown in Fig. 5g, black dashed line.

Finally, using our experimental results in conjunction with our modelling insights, we estimated model parameters for each cell line under consideration (Table I). Recovery times *τ* from averaged FRAP recovery curves (Fig. 3h) and median cortical fractions 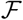 in control and tension-reduced conditions (Fig. 3j) were identified with *F_cor_* and 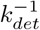 according to Eqn. (2) and (5) (see Materials and Methods). Numerical inversion then gives an estimate of model parameters *k_on_*, *k*_*on*,2_, *k_off_* (0) and *α* (Fig. 6, green analysis pathway). In addition, we used a more elaborate analysis scheme were we employed in addition information about the cortex-to-cytoplasm ratio 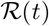 in dependence of timedynamic cortical tension *σ*(*t*) from either cell-squishing experiments or experiments with indirect photo-activation of myosin (Fig. 6, brown analysis pathway). For the simple green analysis scheme, we find significant error bars for estimated model parameter. In particular parameter *k*_*on*,2_ must essentially be regarded as undetermined due to errors in the range of 100% and beyond. Also for parameters *k_off_*(0) and *α*, the green analysis allows only for an order of magnitude estimate. For the brown analysis pathway, error bars are reduced due to the larger set of experimental data fed into the fitting algorithm (Fig. 6). Thus, meaningful estimates can be obtained for all model parameters. Deviations of model parameters between the two analysis pathways can be accounted for by estimated numerical uncertainties. It is noteworthy that our parameter fitting predicts a substantially higher unbinding rate *k_off_*(0) for the GFP-labeled protein as compared to the mKate2-labeled construct which accounts for the lower fraction of cortical Actn4-GFP as compared to Actn4-mKate2 (see Fig. 3c and Fig. 5c). This finding might indicate that unbinding rates are affected by C-terminal tagging of *α*-actinin-4. Most importantly, we find positive slopes of the tension-dependence of the cross-linking time for both *α*-actinin-4 constructs with slope values ranging between 0.4 m/mN and 0.95 m/mN indicating catch bond behaviour of *α*-actinin-4-actin bonds. Within physiological ranges of mean-field cortical tension of 0 — 2mN/m, these slopes put forward a substantial change of *α*-actinin-4 cross-linking time by a factor between 1.8 and 2.9.

**Figure 6.**
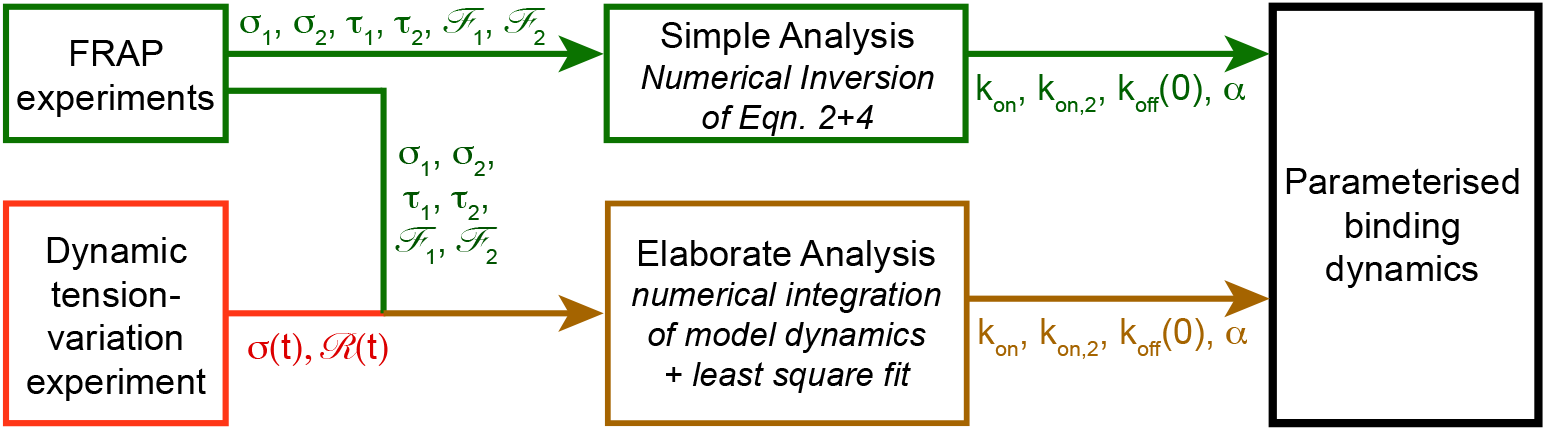
Analysis scheme of experimental data to obtain a parameterised model of cross-linker binding-dynamics. While the green analysis scheme is simpler, the brown analysis scheme provides smaller error bars of determined model parameters, in particular for model parameter *k*_*on*,2_.

We were interested, whether the catch bond tension-dependence of *α*-actinin-4 binding dynamics would be kept in the K255E mutant. The disease-causing mutant K255E of human *α*-actinin-4 has one amino acid of the protein exchanged and exhibits a higher actin-binding affinity than wild-type *α*-actinin-4^9,10,23^. Previous studies reported that this mutant is insensitive to *Ca*^2+^ regulation^9^. Correspondingly, it has been suggested that the K255E mutant of *α*-actinin-4 adopts an open conformation that exposes the additional actin binding site of *α*-actinin-4 outside the ABD and thereby mediates enhanced actin-binding affinity. In turn, observations of a lack of a significant mechanosensitive response in the K255E mutant have been reported^7^.

To investigate the tension-dependence of the K255E mutant further, we transfected HeLa Kyoto cells with a plasmid expressing the GFP-labeled K255E mutant and performed analogous FRAP and AFM measurements (Fig. 7). Consistent with previous observations^7^, we observed fluorescence recoveries of the mutant protein that were substantially slower than that of the wild-type constructs (Fig. 7a). Further, in accordance with previously reported increased actin binding affinity of the K255E mutant^9^, we observed now elevated fractions of cortical *α*-actinin-4 in the mutant as compared to wild-type (Fig. 7c versus Fig. 3c, Fig. 5c). Also different from wild-type *α*-actinin-4, pharmacological reduction of cortical tension did not result in diminished cortical association of the K255E mutant (Fig. 7c). Furthermore, recovery in tension-reduced conditions was slower than in control conditions as reflected by a significantly larger half time of fluorescence recovery (Fig. 7a, control: *t*_1/2_ ≈ 300 s, tension-reduced: *t*_1/2_ ≈ 710 s). Fitting of exponential recovery curves gave recovery time scales τ that did not change significantly upon pharmacological reduction of cortical tension (Fig. 7b). However, the corresponding fitted fraction of recovering fluorescence was significantly lower in tension-reduced conditions (Fig. 7c). This finding puts forward the presence of a subpopulation of *α*-actinin-4 dimers which turns over very slowly, the fraction of which is enhanced if cortical tension is diminished.

**Figure 7.**
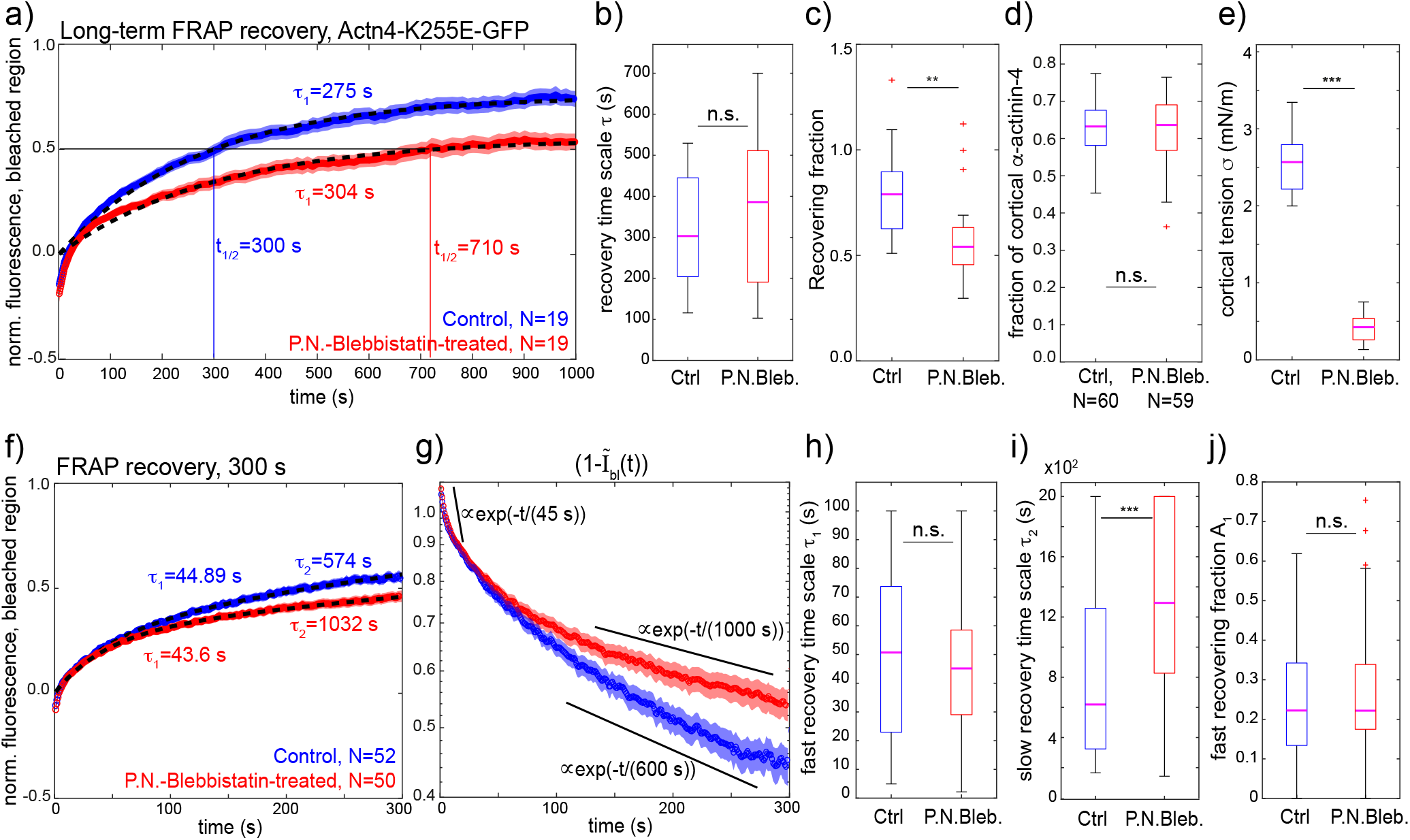
Fluorescence recovery after photobleaching of the K255E mutant of *α*-actinin-4 in the mitotic cortex of HeLa cells. a) Averaged long term recovery of mutant *α*-actinin-4-GFP after photo-bleaching at the mitotic cortex: control (N=19, blue curve), tension-reduced (N=19, red curve, incubated with Para-Nitro-Blebbistatin at 10 *μ*M). Fluorescence intensities were normalized. Recovery curves were averaged and fitted with an exponential recovery with a single time scale (black dashed lines). Red and blue shaded regions indicate the respective standard error of the mean for each condition. b-c) Distribution of parameters from fitting individual recovery curves with an exponential function (same data as panel a): b) Distributions of fit recovery times *τ* (p-value: 0.6). c) Distributions of recovering fractions. d) Estimated fraction of cortical *α*-actinin-4 in control and tension-reduced conditions for an enlarged dataset. Median values of fit recovery time scales are 0.6 in both conditions (control: N=60, P.N.Bleb.: N=59, p-value: 0.3). e) Cortical tensions in control and Blebbistatin-treated conditions corresponding to measurements presented in a. f) Averaged recovery of mutant *α*-actinin-4-GFP during 300 s after photobleaching (large data set). Depicted are normalised fluorescence intensities within the bleached cortex region 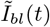. Recovery curves were averaged and fitted with an exponential recovery with two time scales (black dashed lines, see Materials and Methods). Red and blue shaded regions indicate the respective standard error of the mean for each condition. g) Decay of the bleached fraction 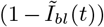 reveals different time scales of relaxations of recovery dynamics (same data as panel f). Note the logarithmic y-axis. h-j) Distribution of parameters from fitting individual recovery curves with a double-exponential function including a fast and a slow recovery time scale corresponding to data. h) Distributions of fitted fast recovery times *τ*_1_ are not significantly different in control and tension-reduced conditions (p-value: 0.56). i) Slow recovery times *τ*_2_ are significantly larger in tension-reduced conditions. j) Distributions of the fraction *A*_1_ of the slow recovering fraction are not significantly changed by cortical tension-reduction (p-value: 0.55). (Measurements are representative for at least two independent experiments. ^*ns*^*p* > 0.05, \**p* < 0.05, \*\**p* < 0.01, \*\*\**p* < 0.001)

To take into account the presence of several time scales in fluorescence recovery, we recorded a larger dataset of mutant *α*-actinin-4 FRAP data focusing on the first 300 s of fluorescence recovery (Fig. 7f-j). We find that fluorescence recovery of averaged curves is well captured by a double-exponential recovery of the form 1 – (*A*_1_ exp(–*t*/τ_1_) + (1 – *A*_1_) exp(–*t*/*τ*_2_))*B*, were *τ*_1_ denotes a fast recovery time scale smaller than 100 s and *τ*_2_ a long recovery time scale > 100 s (Fig. 7a). Systematic fitting of individual recovery curves yields parameter distributions as presented in Fig. 7h-j; we observed no significant differences upon tension reduction in the fast recovering time scale *τ*_1_ and in the fraction of slow fluorescence recovery *A*_1_ (Fig. 7h). By contrast, the slow recovery time scale *τ*_2_ was significantly enlarged upon pharmacological tension reduction (Fig. 7j).

We conjecture that the existence of two recovery time scales is associated to two distinct fluorescent *α*-actinin-4 dimers which are expected to be present in the cell: i) heterodimers that form by dimerization of one (non-fluorescent) endogenous *α*-actinin-4 and one K255E mutant *α*-actinin-4 as well as ii) homodimers that form by dimerization of two fluorescent mutant proteins. In support of this interpretation scheme, we observed that the ratio of slow to fast recovering fractions (1 — *A*_1_)/*A*_1_ increases with the fluorescence intensity of the cortex (Fig. 10, see Materials and Methods for a derivation). Since the K255E mutant of *α*-actinin-4 has significantly higher actin binding affinity^9^, mutant homodimers are expected to exhibit a longer cross-linking time and a slower photobleaching recovery time than heterodimers. Therefore, mutant homodimers are likely associated to recovery time scale *τ*_2_. According to our model, an increase of FRAP recovery time *τ*_2_ upon tension reduction in the actin cortex hints at a slip bond behaviour of mutant homodimers in actin cross-linking (Fig. 7i). In conclusion, our experimental observations of the K255E mutant of *α*-actinin-4 hint not only at a loss of the wild-type catch bond properties of *α*-actinin-4 but also at a trend-reversal of the tension-dependent unbinding rate, i.e. slip bond behaviour.

## IV. DISCUSSION

In this study, we present a new technique that allows to quantify the dependence of actin cross-linker binding on mechanical tension (contractility) in the actin cytoskeletal network of live cells. Therefore, our approach permits to characterise bonds of actin cross-linkers to be of the slip or catch bond type (or tension-insensitive) in a physiological range of active tensions in the cellular cortex. As part of our assay, we performed FRAP experiments on fluorescently-labeled cross-linker proteins in round mitotic cells in combination with AFM-based cell confinement and a meanfield mathematical model of cross-linker binding dynamics. We applied this technique in an exemplary manner to characterise the binding dynamics of the actin cross-linker *α*-actinin-4, which is the most abundant actin cross-linker in HeLa cells^24^.

In terms of our mathematical model, we could connect cross-linking lifetime of a cross-linker molecule to its overall residence time at the cortex. We showed, that there is a qualitative correspondence between the tension-dependence of cross-linking lifetime and cortical residence time: if cross-linking lifetime increases in dependence of cortical tension, also cortical residence time increases (catch bond scenario). On the other hand, if cross-linking time decreases in dependence of tension, our model predicts a corresponding decrease of cortical residence time (slip bond scenario). However, the respective magnitude of the slopes may differ between cross-linking lifetime and cortical residence time (Fig. 2a-c, rightmost panels). We recorded experimental data of FRAP-derived cortical residence times of *α*-actinin-4 in dependence of cortical tension and used our theoretical insight to derive associated actin-binding and unbinding rates of *α*-actinin-4 (Table I). In conclusion, we find that estimated cross-linking lifetimes and cross-linking concentrations of wild-type *α*-actinin-4 increase by a factor 2-3 within a physiological tension range of cortical tension of 0.1 – 2 mN/m^11,12,15,25^, rendering *α*-actinin-4 a catch bond cross-linker in cells (Fig. 2b,c,e). By contrast, for the K255E mutant of *α*-actinin-4, signatures of catch bond binding as predicted by our model were missing such as higher cortical association and longer cortical residence times at higher cortical tension. In fact, the observed phenotype of slower FRAP recovery at reduced cortical tension points at a slip bond behaviour of the mutant protein.

Cross-link concentration is a parameter that sensitively regulates the stiffness of cross-linked biopolymer networks such as the actin cytoskeleton^26–28^. Furthermore, cross-linking lifetime was shown to introduce characteristic time scales into the viscoelastic mechanics of a cross-linked biopolymer network since cross-linker unbinding gives rise to local stress-releases^14,29^. Therefore, tension-regulated manipulation of cross-linking lifetimes and cross-link concentration represents an intriguing mechanism for the cell to tune its cytoskeletal structure, stiffness and mechanical relaxation times through active mechanical tension. In particular, understanding and quantifying tension-dependence of actin cross-linker binding is key for the understanding of nonlinear material properties of the cytoskeleton^30–32^. The approach presented here now allows to systematically study to which extent this mechanism is implemented in cells.

## Appendix Materials and Methods

### 1. Cell culture

We cultured HeLa Kyoto cells expressing a green-fluorescent or red-fluorescent, C-terminally tagged *α*-actinin-4 construct in DMEM (PN:31966-021, life technologies) supplemented with 10% (vol/vol) fetal bovine serum, 100 *μ*g/ml penicillin, 100 *μ*g/ml streptomycin and 0.5 *μ*g/ml geneticin (all Invitrogen) at 37°C with 5% CO_2_. One day prior to the measurement, 10.000 cells were seeded into a silicon cultivation chamber (0.56 cm^2^, from ibidi 12 well chamber) that was placed in a 35 mm cell culture dish (fluorodish FD35-100, glass bottom) such that a confluency of ≈ 30% is reached at the day of measurement.

For AFM experiments, medium was changed to DMEM (PN:12800-017, Invitrogen) with 4 mM NaHCO_3_ buffered with 20 mM HEPES/NaOH pH 7.2. Mitotic arrest of cells was achieved by addition of S-trityl-L-cysteine (STC, Sigma) two to eight hours before the experiment at a concentration of 2 *μ*M. This allowed conservation of cell mechanical properties during measurement times of up to 30 min for one cell^33^. Cells in mitotic arrest were identified by their shape and visibility of condensed DNA in transmitted light images. Pharmacological agents (-)-Blebbistatin (sigma, B0560-1MG), Para-Nitro-Blebbistatin (Optopharm, DR-N-111) or Y-27632 (Cayman Chemical, Cay10005583-1) were added at least 15 min prior to measurement to indicated concentrations. As opposed to Blebbistatin^20^, Para-Nitro-Blebbistatin is optically stable and allows cell imaging with blue light without activity changes of the drug.

### 2. Cell lines

Within this study three different HeLa cell lines were used expressing either fluorescently labeled murine Actn4 from BACs (see Fig. 3 and 5) or GFP-labeled human Actn4 with a point mutation (K255E) from a plasmid. The K255E mutant was cloned into the plasmid backbone pLGC as described in^34^. Hela Kyoto cells were stably transfected with this plasmid using Lipofectamine 2000 (Invitrogen, 11668030) according to the protocol of the manufacturer.

BACs with Actn4 constructs were prepared as follows: BACs harboring mouse Actn4 (RP23-375C17) were modified by recombineering as described^35^. Briefly, using electroporation, both, a plasmid pSC101 carrying two recombinases and gene-specific purified EGFP- or mKate2-tagging cassettes were introduced into the E. coli strains containing the parental BAC vectors. For this purpose, 50 nucleotides-homology arms that matches the C-termini of both target genes, respectively, were attached to the BAC tagging cassettes using PCR. Precise incorporation of the tagging cassette was confirmed by PCR and sequencing. Next, tagged BACs were isolated from bacteria using the Nucleobond PC100 kit (Macherey-Nagel, Germany). Subsequently, HeLa Kyoto cells were transfected using Effectene (Qiagen) and cultivated in selection media containing 400 *μ*g/ml geneticin (G418, Invitrogen). Finally, pools of HeLa cells stably expressing Actn4-EGFP were analyzed by western blot and immunofluorescence using an anti-GFP antibody (11814460001, Roche) or to verify correct protein size and localization of the tagged transgene.

### 3. Confocal imaging in combination with AFM

The experimental set-up consisted of an AFM (Nanowizard I, JPK Instruments) mounted on a Zeiss LSM700 laser scanning confocal system of the CMCB light microscopy facility. For imaging, we used a 20x objective (Zeiss, Plan Apochromat, NA=0.80) and a 555 nm laser line for excitation of mKate2 fluorescence at laser powers between or a 488 nm laser line for excitation of GFP both at laser powers between 0.3 – 1 %. During measurements, cell culture dishes were kept in a petri dish heater (JPK instruments) at 37°C. On every measurement day, the spring constant of the cantilever was calibrated using the thermal noise analysis (built-in software, JPK). Cantilevers were tipless, 200 – 350 *μ*m long, 35 *μ*m wide, 2 *μ*m thick (NSC12/tipless/noAl or CSC37/tipless/noAl, Mikromasch) with nominal force constants between 0.3 and 0.8 N/m. Cantilevers were modified with wedges to correct for the 10° cantilever tilt consisting of UV curing adhesive (Norland 63)^36^.

Prior to cell confinement, the AFM cantilever was lowered to the dish bottom near the cell until it came into contact with the glass surface and then retracted to ≈ 14.5 *μ*m above the surface. Thereafter, the free cantilever was moved over the cell. The confocal plane of scanning was adjusted to coincide with the equatorial plane of the cell that exhibits the largest cross-sectional area. To increase fluorescence intensity, the aperture of the pinhole was set to 3 Airy units corresponding to an optical section thickness of 6 *μ*m.

Stationary state AFM forces were used to calculate cortical tension as described previously^12,17^. Briefly, as part of this analysis, cell height and cross-sectional area of the cell in the equatorial plane were used to estimate cell volume and other geometrical parameters (contact area, mean curvature) as described in^12^.

During the entire experiment, the force acting on the cantilever was continuously recorded at a frequency of 10 Hz. The height of the confined cell was computed as the difference between the height that the cantilever was raised from the dish surface and lowered onto the cell plus the height of spikes at the rim of the wedge (due to imperfections in the manufacturing process^36^) and the force induced deflection of the cantilever.

### 4. Photo-bleaching experiments

AFM cell confinement of a cell in mitotic arrest was realised as described in the previous section at a height of ≈ 14.5 *μ*m (retracted AFM cantilever) and recording of AFM force and cantilever height was started. Afterwards, confocal imaging of the chosen cell was set up. For every cell, we scanned a square region of 512 × 512 pixels (side length between 30 – 50 *μ*m) that covered the equatorial cell cross-section and partly its surrounding. We recorded a time series at time intervals between 1.5 and 4s. After imaging of 4-10 initial frames, a small rectangular region of ≈ 2 × 20 *μ*m^2^ of the cell cortex was bleached in 3 iterations with the 555 nm laser at a laser power of 70% and a pixel dwell time 2.55 *μ*s (see Fig. 8a, green rectangle depicts bleached area). After bleaching a time interval of 150 – 250s was recorded until fluorescence in the bleached area plateaued.

**Figure 8.**
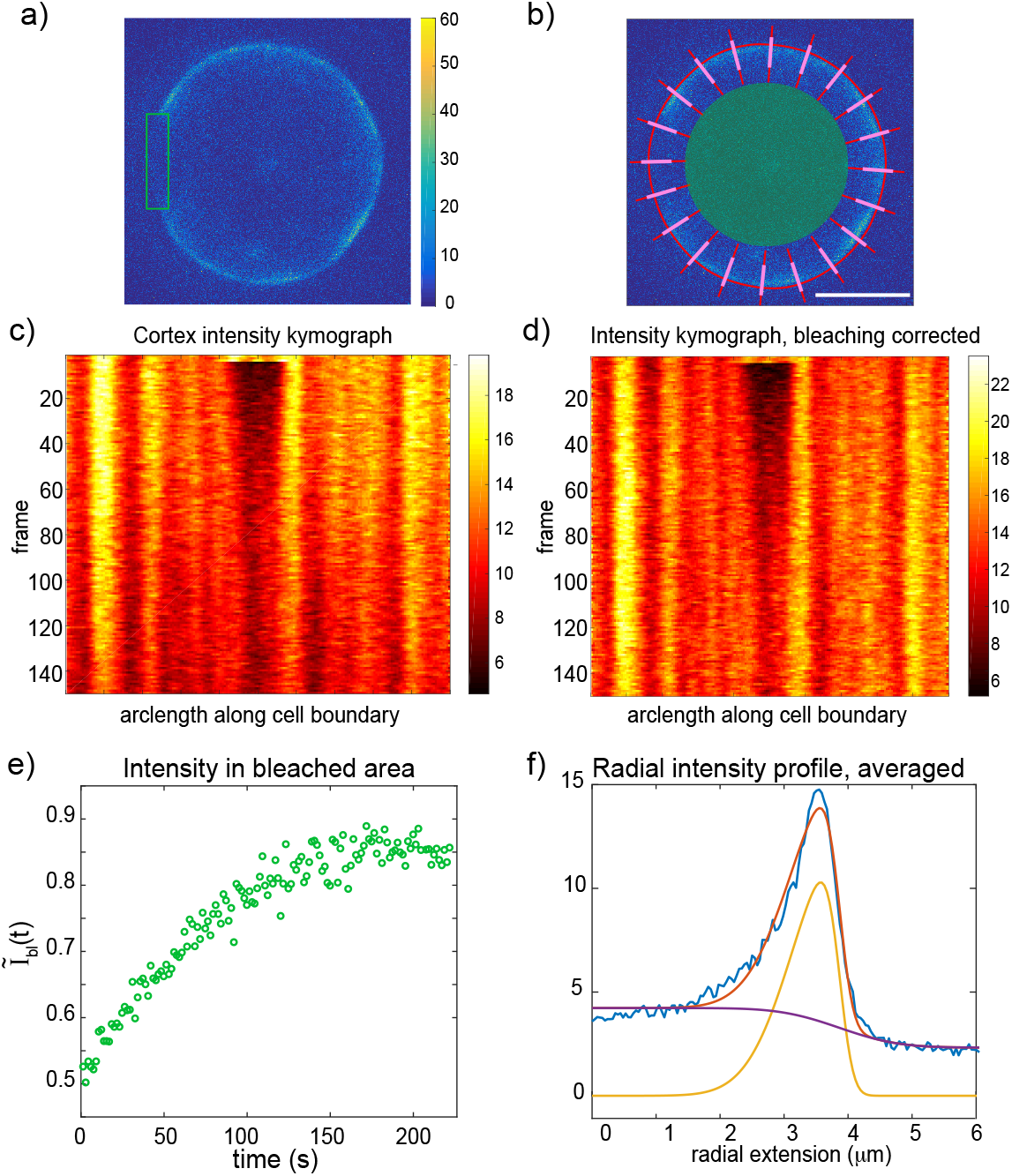
Exemplary analysis of photo-bleaching recovery in a mitotic HeLa cell with mKate2-labeled *α*-actinin-4. a) Confocal Micrograph of the equatorial cell cross-section right after photo-bleaching. The bleached region is indicated by the green rectangle. b) Same micrograph as in panel a showing now elements for image analysis generated by our analysis code: the identified cell boundary (red cell outline), orthogonal lines to the cell boundary (red and rosa lines, only every tenth line plotted for illustration purposes) and an inner circular cytoplasmic region (green disk) that extends up to 70% of the cell radius. c) Kymograph showing cortical fluorescence over time. Kymographs were generated by plotting integrated fluorescence intensities for each orthogonal rosa line (see panel b) for every time point. d) Kymograph as in panel c but with bleaching correction. Notably, fluorescence intensities outside the bleached region now remain constant on average. e) Averaged and normalised fluorescence intensity over time in the bleached region of the cortex (see panel c). Intensity values were corrected for photo-bleaching. f) Fluorescence intensity profile averaged along the cell circumference before photo-bleaching (blue curve). A smoother version of the fluorescence profile is obtained by fitting function (A.1) (red curve), which consists of a cortical skewed Gaussian (yellow curve) and a contribution that captures cytoplasmic and background fluorescence (violet curve).

#### Image analysis of photo-bleaching experiments

For image analysis of confocal time series’, the cell boundary was determined by a matlab custom-code (see Fig. 8b for an example cell, the cell boundary is marked in red). Along this cell boundary, 200 orthogonal, equidistant lines were determined extending 4μm to the cell interior and 2*μ*m into the exterior (see Fig. 8b, red lines orthogonal to cell boundary, only every 10th line is plotted out of 200).

To obtain fluorescence recovery curves from the bleached region of the cell cortex, we determined the averaged intensity along the cortex circumference by averaging fluorescence intensity values along 200 shorter radial lines (rosa lines in Fig. 8b, extending 2.4*μ*m to the interior, 0.8*μ*m to the exterior). Plotting average intensities along the cortex circumference over time allows to generate a kymograph of cortex fluorescence intensity, see Fig. 8c.

We determined the bleached area of the cortex, by identifying the region with more than 40% intensity change from the frame before to the frame after bleaching. The intensity average along rosa, radial lines within the bleached cortical area *I_bl_* was calculated for every time point. Furthermore, we determined the averaged intensity in the unbleached cortical region *I_un_* over time *t*. To estimate the rate of bleaching *k_bl_*, we fitted the function *I_un_*(*t*) by an exponential decay *A*exp(–*k_bl_t*). Correcting now intensity values along the cortex by multiplying it with the factor exp(*k_bl_t*), we may plot a new kymograph, where bleaching has been corrected (Fig. 8d), such that the average intensity in the unbleached area remains constant.

The normalized and bleaching-corrected intensity 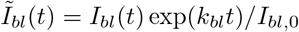 was then in the following used to analyse fluorescence recovery (see Fig. 8e), where *I*_*bl*,0_ denotes the intensity bleached region before bleaching. Recovery curves 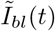 were fit by a an exponential rise

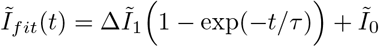

in terms of a least-squares fit. The fit function takes the value 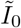 at *t* = 0 but converges to 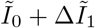 at long times. Recovery is captured by a characteristic time scale *τ*. The first 5 s of the recovery were excluded from fitting, since in this time regime, it was apparent that recovery had a major contribution from cytoplasmic diffusion into the cytoplasmic region next to the bleached cortex area. Renormalized intensities 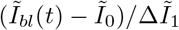 where then averaged for all cells measured to obtain Fig. 3d, h and 5a, main text.

#### Estimating the fraction 𝓕 of cortical fluorescent protein

Fluorescence intensities of radial lines at the cell periphery (red lines in Fig. 8b) were averaged for all pre-bleaching frames to obtain an average intensity profile in radial direction along the cortex (see Fig. 8f, blue curve). This intensity profile along the red orthogonal line is then fitted by the function

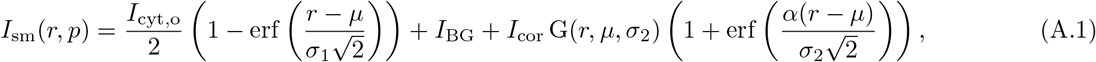

with fit parameters *p* = {*μ*, *σ*_1_, *σ*_2_, *I*_cyt,o_, *I*_cor_, *I*_BG_, *α*}, to obtain a smoothed variant *I*_sm_(*r,p*) of the intensity profile along the radial coordinate. A respective exemplary fit of Eq. (A.1) is shown in Fig. 8f, orange curve. Here, erf(*r*) denotes the error function and G(*r, μ, σ*_2_) denotes a Gaussian function with mean value *μ* and standard deviation *σ*_2_. The term 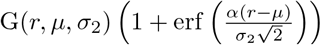 is a corresponding skewed Gaussian function with skewness characterized by parameter *α*. The parameters *I*_cyt,o_ and *I*_cor_ quantify the amplitude of cytoplasmic versus cortical fluorescence, while *σ*_1_ and *σ*_2_ quantify the smearing out of the cytoplasmic or cortical fluorescence distribution, respectively. *I_BG_* captures the value of background intensity, while *μ* is the coordinate for the location of the cell boundary.

The skewed Gaussian contribution in the fitted intensity profile (Fig. 8f, yellow curve) is interpreted as fluorescence contribution from the cortical proteins, while the remaining part (Fig. 8f, violet curve) is interpreted as a sum of cytoplasmic fluorescence in the outer rim of the cell *I_cyt,o_* and background fluorescence *I_BG_*. The skewed Gaussian was then integrated along the radial direction to obtain *I*_2*D,cortex*_ as a measure for the 2D concentration of fluorescent protein at the cortex. The integrand was then multiplied with the overall cell surface area *A_cell_* to an intensity value that is proportional to the overall number of fluorescent proteins *N_cor_* at the cortex. To estimate the cytoplasmic concentration of fluorescent protein, we determine the average fluorescence intensity in the inner part of the crosssection *I_cyt,i_* (Fig. 8b, green disk extending to 70% of the cell radius) and in the outer cytoplasmic region *I_cyt,o_*. *I_cyt,o_* is the innermost value of *I_sm_* of a radial line profile (Fig. 8b, red lines). To obtain an overall intensity average of the cytoplasmic fluorescence, we calculate *I_cyt_* = (0.7^2^ *I_cyt,i_*¿ + (1^2^ – 0.7^2^)*I_cyt,o_*) – *I_wedge_*, where *I_wedge_* is the background intensity from the wedge, which we estimate as 0.46 × *I_BG_* (see Section below and Fig. 9). *I_cyt_* × *V_cell_* is then used as a measure for the number of fluorescent proteins in the cytoplasm *N_cyt_*, where *V_cell_* is the estimated volume of the cell. Therefore, we approximate the fraction of cortical fluorescent protein 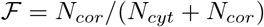 as (*I*_2*D,cort*_*A_cell_*)/(*I_cyt_V_cell_* + *I*_2*D,cort*_A_cell_).

**Figure 9.**
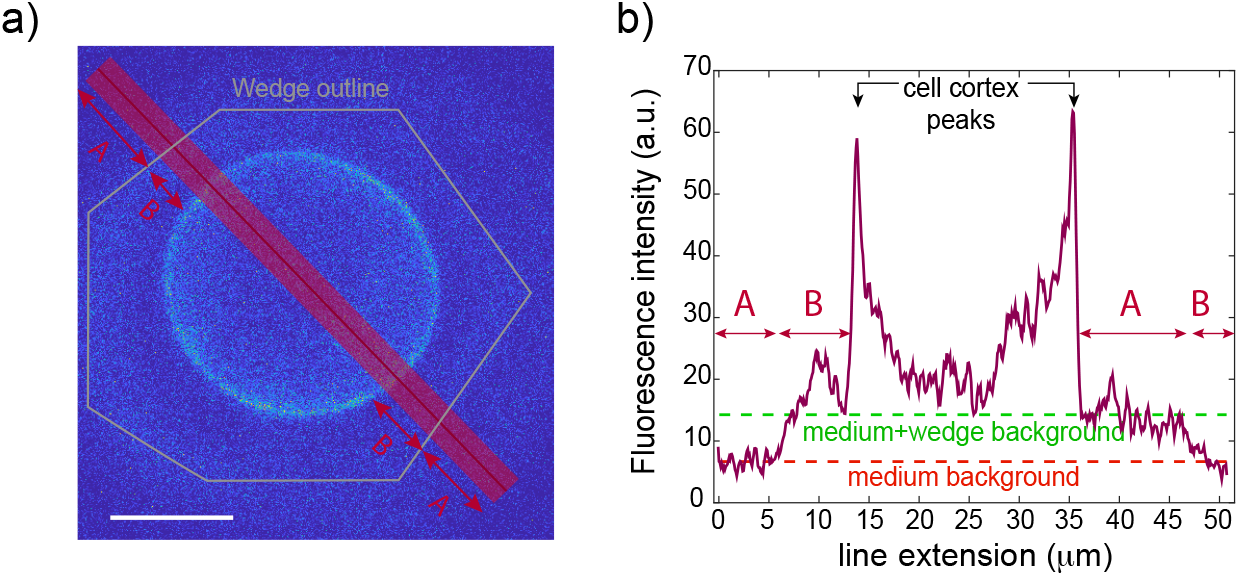
Quantification of background fluorescence from the cantilever wedge. a) Exemplary picture of fluorescence profile of the equatorial cross-section of the cell with surrounding. The cantilever wedge (indicated by the grey outline) is a source of autofluorescence. b) A line scan across the cell shows the fluorescence profiles in three different areas: outside the cantilever wedge area (A), underneath the cantilever wedge but outside the cell (B), underneath the cantilever wedge inside the cell (central region). Inside area A, fluorescence stems from autofluorescence of the medium. Inside area B, fluorescence emanates from autofluorescence of the medium and the wedge. Inside the central region, fluorescence stems from fluorescence of cellular components and autofluorescence of the wedge.

Due to uncertainty in the background fluorescence, we estimate the relative error of *I*cyt** to be ≈ 20%. Due to error propagation, we estimate a corresponding relative error of up to 10% of 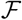.

As previously described^12^, we estimated the cell volume *V_cell_* and the cell surface area *A_cell_* as

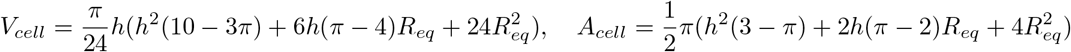

which can be derived by regarding the cell as an axisymmetric body of revolution and approximating the profile of the free cell contour by a semi-circle with radius *h*/2. Here, *R_eq_* is the equatorial radius of the confined cell (obtained from imaging) at confinement height *h* (obtained as AFM readout).

### 5. Blue light inactivation experiments

After moving the cantilever over the cell and a subsequent initial equilibration time of at least 200 s, we started to record a time series of confocal images with the focal plane coinciding with the equatorial plane of the cell choosing time intervals between 5 — 10 s. After imaging between 5-10 initial frames, the frame was laser-scanned with blue light (488 nm, diode laser of Zeiss LSM 700) in 10 iterations at a laser power of 8%, speed of 10 (pixel dwell time 1.27 *μ*s) to inactivate Blebbistatin in the cell^20^. To monitor the concentration of cortex-bound fluorescent protein during photo-inactivation of Blebbistatin, we determined the averaged intensity at the cell boundary *I_bound_* by integrating fluorescence intensities over all 200 short radial lines at the cortex (rosa lines in Fig. 8b, extending 2.4*μ*m to the interior, 0.8*μ*m to the exterior). Here, averaging was performed along the line and over all 200 lines along the circumference. To quantify the cytoplasmic concentration of fluorescent protein, we determined the average fluorescence intensity in the inner part of the cross-section *I_cyt,i_* (Fig. 8b, green disk extending to 70% of the cell radius). To estimate the ratio of cortex-bound to cytoplasmic fluorophore concentration in Fig. 3 and 5, we calculate the ratio (*I_bound_*–*I_wedge_*)/(*I_cyt,i_* – *I_wedge_*) where *I_wedge_* is the background intensity from the wedge, which we estimate as 0.46 × *I_BG_* (see Section below and Fig. 9b and following paragraph).

#### Fluorescence background analysis

For image analysis, we quantified contributions to background fluorescence by plotting the intensity profile along a straight line spanning the cell diameter and its neighborhood (Fig. 9a). We identified two contributions to the background fluorescence: i) auto-fluorescence of the wedge and ii) auto-fluorescence of the medium. Estimating average fluorescence values in regions A and B, respectively, as indicated in Fig. 9a,b, we quantified background fluorescence in the equatorial plane of the cell at a cantilever height of ≈ 14 *μ*m for six cells. We obtained an average value of 6.7 ± 1.7 (error indicates the standard deviation) in the region outside of the wedge (region A), while we quantified an average value 11.2±3.4 in the region underneath the wedge (region B). This indicates that the background fluorescence from the wedge adds ≈ 46 % to the overall fluorescence background, while the remaining background is constituted by the auto-fluorescence of the medium.

### 6. Cell-squishing experiments

After moving the retracted cantilever over the cell at a height of *h* ≈ 14.5 *μ*m, we let the cell equilibrate for an initial equilibration time of at least 200 s. Then, a time series of confocal imaging was started while the cell was still confined by a retracted cantilever. After imaging between 2-10 initial frames, cell-squishing was started by lowering the cantilever at a speed of 0.5 *μ*m/s down to a nominal height between 9.5 and 11.5μm. Imaging was performed at time intervals between 1.5 – 5s. Before the start of the imaging time series, the focal plane was adjusted to coincide with the equatorial plane after cell-squishing was finalized.

We registered AFM force curves and imaging time series of the cell in time by setting the following events to be at time zero: i) the peak of the AFM force curve and ii) the first frame in the imaging series where the cell crosssection does not continue to grow. Images recorded before this time point were not used for analysis. For averaging cortex-to-cytoplasm ratios from several cells, the measured ratio 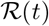 was normalised for each cell through division by smoothed 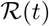 curve evaluated at time t = 150 s. In this way, averaged normalised cortex-to-cytoplasm ratios converge to one in the long time limit (see Fig. 4c). To test the significance of the peak at time *t* = 30s after uniaxial compression, we applied a T-test to the distribution of normalised intensity ratios for ten cells and found that the mean value is significantly different from 1 (p-value: 0.041, mean: 1.04).

### 7. Model parameter determination

To determine model parameters for different *α*-actinin4 constructs, we used for each construct experimentally measured quantities 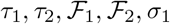 and *σ*_2_; *τ*_1_ and *τ*_2_ are measured recovery times of averaged FRAP recoveries in condition 1 (control) or condition 2 (tension-reduced), respectively (Fig. 3d,h, Fig. 5a). Correspondingly, 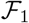 and 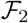 denote the measured median fractions of cortical protein in both conditions as depicted in Fig. 3f,j and Fig. 5c. *σ*_*act*1_ and *σ*_*act*,2_ denote respective measured median values of cortical tension as depicted in Fig. 3g,k and Fig. 5d.

For mutant protein Actn4-K255E-GFP, we determined model parameters as presented in Table I by directly solving the following equations numerically with respect to *k_on_*, *k*_*on*,2_, *k_off_*(0) and *α*:

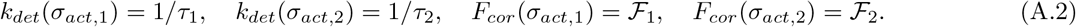

Here, *k_det_*(*σ*_*act*1/2_) and *F_cor_*(*σ*_*act*1/2_) are given by Eq. (5) and (2), respectively. This corresponds to the green analysis pathway as depicted in Fig. 6.

For cell lines with BAC constructs (Actn4-EGFP, Actn4-mKate2), we determined model parameters according to the brown pathway as depicted in Fig. 6. There, we used in addition to the above mentioned experimental quantities 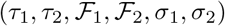 the output from cell-measurements with dynamic tension variation (cell-squishing or indirect photo-activation). This output includes times series of cortex-to-cytoplasm ratio of cross-linkers 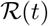 and cortical tension *σ*(*t*). The experimental quantity 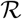 corresponds to the model quantity *R* as given in Eq. 3. Using all of these experimentally measured quantities, we performed a least squares fit to obtain model parameters that provide the best match. A nonlinear least squares fit was performed using matlab and the command lsqnonlin. The least squares fit was minimising the following functional

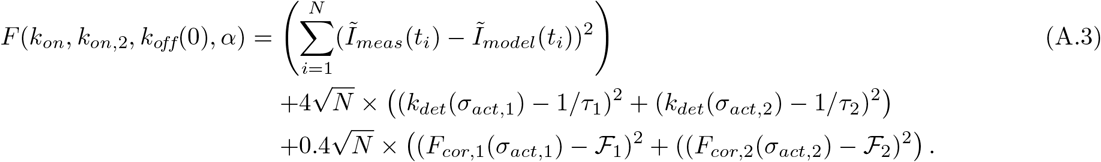

Here, 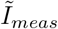 denotes the averaged normalised fluorescence intensities from measurements as depicted in Fig. 3d and Fig. 5h, while 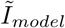 where corresponding model predictions of normalised intensity.

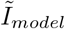 was obtained from numerical integration of defining Eqn. (2) in matlab using a Euler forward algorithm with time step Δ*t* = 0.1 s in matlab. 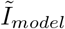 was calculated (*c_sb_*(*t*) + *c_d_*(*t*))/*c_cyt_*(*t*) × *C_cyt_*(*t_end_*)/(*c_sb_*(*t_end_*) + *c_cl_*(*t_end_*)), where *t_end_* is the biggest time argument under consideration.

For both, the green and the brown analysis pathway, errors of estimated model parameters were determined by calculating derivatives of all model parameters with respect to each measured variable. The error of the model parameter was then estimated by the variance formula of error propagation. Accordingly, the error of *k_on_* was calculated as

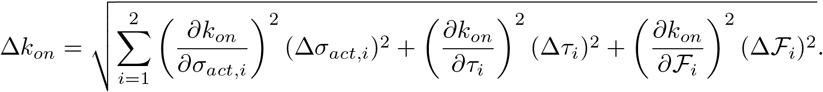

For all other model parameters, the error estimate was performed analogously. The relative errors of *σ*_*act*,1_, *σ*_*act*,2_, *τ*_1_ and *τ*_2_ were estimated as 5%, while the relative errors of 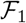 and 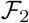 were estimated as 10%. We find that the brown analysis pathway reduces error bars in particular for the attachment parameter *k*_*on*,2_ (see Table I).

### 8. Numerical integration of the model

Simulations of model Eqn. (2) presented in Fig. 2 were obtained by Mathematica using the command NDSolve.

For data fitting as presented in Fig. 3h and Fig. 5g, numerically integrated solutions of our model where fitted to measured data by a nonlinear least squares fit using matlab and the command lsqnonlin (see previous section).

For the simulation of cell-squishing experiments, the anticipated squishing-induced increase of cell surface area was taken into account. Experimentally, we identified a typical cell volume of ≈ 5000 *μ*m^3^ of mitotic HeLa cells. Therefore, using the assumption of cellular droplet shapes^11^, we can predict cellular surface area increase due to squishing to be between ≈ 6.5 – 15% for an initial cell height of 14.5 *μ*m and final heights between 9.5 – 11.5*μ*m. For the numerical integration of our model, we assumed an average increase of this ratio by 11%. For simulations of cell-squishing experiments and corresponding data fitting, surface areas were for simplicity approximated to increase linearly from the starting point of cantilever lowering until cantilever halt. Surface area increase of cells through squishing can account for the observed elevated equilibrium intensity of *α*-actinin-4 after squishing, see Fig. 3h.

### 9. Analysis of fluorescence recovery for the K255E mutant

Recovery curves of the K255E mutant were not fit well by an exponential recovery with a single time scale. Instead, fluorescence recovery of averaged curves were well captured by a double-exponential recovery of the form 1 – (*A*_1_ exp(–*t*/*τ*_1_) + (1 – *A*_1_) exp(–*t*/*τ*_2_))*B*, were *τ*_1_ denotes a fast recovery time scale smaller than 100 s and *τ*_2_ a long recovery time scale > 100 s. Since mutant *α*-actinin-4 has significantly higher actin binding affinity^9^, mutant homodimers are expected to exhibit a longer cross-linking time and a slower photobleaching recovery time than heterodimers consisting of a mutant and a wild-type protein. In fact, in the framework of our model, heterodimers will exhibit an effective unbinding rate 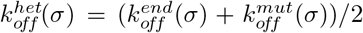 where 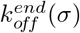 and 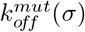 are the un binding rates of one ABD in the cross-linking state for endogenous and mutant *α*-actinin-4, respectively. Therefore, the tension-dependent unbinding rate 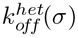 of heterodimers is a mixture of the tension-dependence of endogenous and mutant protein. It is noteworthy that in the limit case of small or negligible tension-dependence of mutant unbinding, heterodimers would effectively behave as catch bond cross-linkers. Further, it is possible that the two tension-dependences of wild-type and mutant bond effectively cancel each other out. The observation of no significant changes of the fast FRAP recovery time *τ*_1_ upon tension reduction (see Fig. 7h) is in support of this latter scenario. If the fast recovery time *τ*_1_ is associated to heterodimer recovery and the slow recovery time *τ*_2_ to homodimer recovery, the ratio (1 – *A*_1_)/*A*_1_ is expected to increase with the fluorescence intensity of the cell by the following proportionalities:

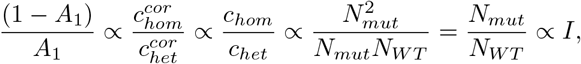

where 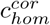 and 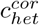 are the cortical concentration of homodimers or heterodimers, *c_hom_* and *c_het_* are the cell-averaged concentrations of homo- and heterodimers, *N_mut_* and *N_WT_* are the number of mutant and wild-type *α*-actinin-4 proteins in the cell and *I* is the fluorescence intensity of the cell. This proportionality is confirmed by a significant positive correlation between 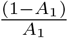 and absolute cortical fluorescence intensities of cells (Fig. 10).

**Figure 10.**
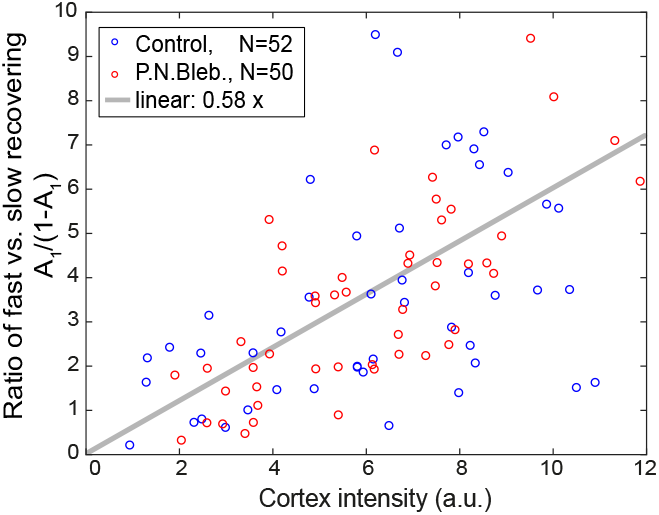
Ratio of slow versus fast recovering fractions versus absolute cortical fluorescence intensities (same data as panel Fig. 7f). The Pearson correlation coefficient was determined as 0.43 with a p-value of ≈ 10^−5^ (data points of both conditions were pooled).

## ACKNOWLEDGMENTS

We thank William Brieher for provision of plasmids expressing ACTN4^K255E^-GFP. Further, we thank Anthony Hyman for provision of transgenic HeLa cell lines. In addition, we thank the CMCB light microscopy facility for excellent support. EFF thanks for financial support from the DFG, project FI 2260/4-1. KH and EFF were further supported by the Deutsche Forschungsgemeinschaft (DFG, German Research Foundation) under Germany’s Excellence Strategy, EXC-2068-390729961, Cluster of Excellence Physics of Life of TU Dresden.

## AUTHOR CONTRIBUTIONS

E.F.F. designed the research. K.H. and E.F.F. performed experiments and data analysis. L.S. and E.F.F. developed matlab scripts for image and data analysis. E.F.F. developed the theoretical underpinning for data analysis and performed numerical simulations. I.P. generated cell lines from BACs. E.F.F. and K.H. wrote the manuscript.

